# Patterning, regulation, and role of FoxO/DAF-16 in the early embryo

**DOI:** 10.1101/2024.05.13.594029

**Authors:** Michael S. Mauro, Sophia L. Martin, Julien Dumont, Mimi Shirasu-Hiza, Julie C. Canman

## Abstract

Insulin resistance and diabetes are associated with many health issues including higher rates of birth defects and miscarriage during pregnancy. Because insulin resistance and diabetes are both associated with obesity, which also affects fertility, the role of insulin signaling itself in embryo development is not well understood. A key downstream target of the insulin/insulin-like growth factor signaling (IIS) pathway is the forkhead family transcription factor FoxO (DAF-16 in *C. elegans*). Here, we used quantitative live imaging to measure the patterning of endogenously tagged FoxO/DAF-16 in the early worm embryo. In 2-4-cell stage embryos, FoxO/DAF-16 initially localized uniformly to all cell nuclei, then became dramatically enriched in germ precursor cell nuclei beginning at the 8-cell stage. This nuclear enrichment in early germ precursor cells required germ fate specification, PI3K (AGE-1)- and PTEN (DAF-18)-mediated phospholipid regulation, and the deubiquitylase USP7 (MATH-33), yet was unexpectedly insulin receptor (DAF-2)- and AKT-independent. Functional analysis revealed that FoxO/DAF-16 acts as a cell cycle pacer for early cleavage divisions–without FoxO/DAF-16 cell cycles were shorter than in controls, especially in germ lineage cells. These results reveal the germ lineage specific patterning, upstream regulation, and cell cycle role for FoxO/DAF-16 during early *C. elegans* embryogenesis.

## Introduction

The insulin/insulin-like growth factor signaling (IIS) pathway plays a critical role in metabolism, immunity, environmental stress resistance, reproduction, and aging. When the IIS pathway is overstimulated, insulin resistance can develop into type 2 diabetes, which has become extremely prevalent throughout the world and in particular the United States, where an estimated ∼40 million people (∼10% of the population) are affected (CDC, https://www.cdc.gov/diabetes/data/statistics-report/index.html#anchor_86793). Insulin resistance and diabetes lead to a variety of adverse health effects including cardiovascular disease, blindness, kidney disease, and fertility issues in both men and women (Deshpande et al., 2008; Jangir and Jain, 2014; La Vignera et al., 2012; Livshits and Seidman, 2009; Thong et al., 2020). Insulin resistance and diabetes during pregnancy can also be detrimental to embryonic and fetal development, leading to an increased rate of miscarriage and birth defects (Cai et al., 2022; Clausen et al., 2005; Correa et al., 2008; Craig et al., 2002; Tian et al., 2007). However, given the tight relationship between insulin signaling and obesity, which also impacts fertility, it has been difficult to identify the exact roles of the IIS pathway in pregnancy and early embryonic development.

In *C. elegans*, the first screen to identify a key role for the IIS pathway was a genetic screen to uncover genes involved in dauer formation, a developmentally arrested larval stage that occurs upon starvation or overcrowding (Albert et al., 1981). More recently, the worm IIS pathway has also been implicated in regulating reproduction. (Das and Arur, 2017; Hubbard, 2011; Luo et al., 2010; Michaelson et al., 2010; Qin and Hubbard, 2015; Tissenbaum and Ruvkun, 1998) Specifically, the worm IIS pathway controls germline proliferation (Michaelson et al., 2010), oocyte quality (Luo et al., 2010; Templeman et al., 2018), and oocyte maturation (Lopez et al., 2013). While much attention has been paid to the role of the IIS pathway in larvae and adults, less is known about how the pathway functions during embryonic development.

DAF-16 is the sole FoxO (forkhead family transcription factor class O, hereafter FoxO^DAF-16^) ortholog in *C. elegans* and a critical downstream target of the IIS pathway (**Figure 1A**) (Furuyama et al., 2000; Murphy and Hu, 2013). When conditions are favorable (*e.g.*, in food abundance), insulin binds to DAF-2/insulin receptor (hereafter InR^DAF-2^), activating the phosphatidylinositol (PI)-3 kinase (AGE-1 in worms, hereafter PI3K^AGE-1^), leading to downstream activation of PDK-1 (3-phospholipid-dependent kinase), AKT-1/AKT-2/SGK-1, and phosphorylation of FoxO^DAF-16^, preventing it from entering the nucleus (**Figure 1A**) (for review see (Murphy and Hu, 2013; Tissenbaum, 2018)). When conditions are unfavorable and InR^DAF-2^ is inactive (*e.g.*, upon starvation), the lipid phosphatase DAF-18/PTEN, hereafter PTEN^DAF-18^, converts PIP_3_ to PIP_2_, deactivating PDK-1 and AKT family kinases and allowing FoxO^DAF-16^ entry into the nucleus to alter gene expression (**Figure 1A**). (Tissenbaum, 2018) FoxO^DAF-16^ is also implicated in a variety of other metabolic pathways such TOR (target of rapamycin), AMP kinase, and JNK (c-Jun N-terminal kinase) pathways in response to nutrient changes and other environmental ques (Landis and Murphy, 2010; Sun et al., 2017). Through these pathways, FoxO^DAF-16^ plays a transcriptional role in regulating cell cycle timing, tumor suppression, and establishment and maintenance of the germline (Baugh and Sternberg, 2006; Curran et al., 2009; Hornsveld et al., 2021; McElwee et al., 2003; Pinkston-Gosse and Kenyon, 2007; Qi et al., 2017; Yang et al., 2013).

**Figure 1:**
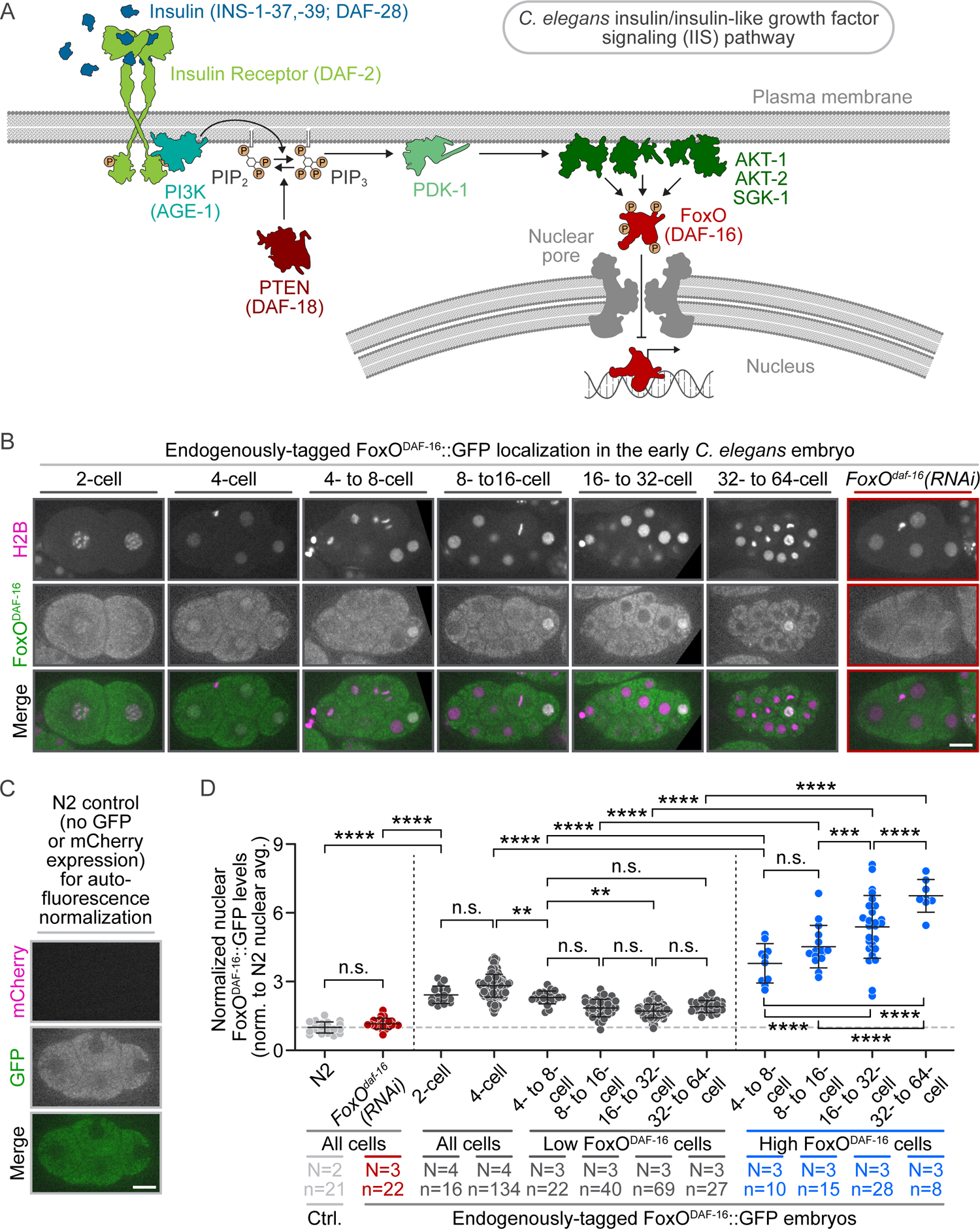
FoxODAF^-16^ is expressed in the early *C. elegans* embryo and becomes enriched in posterior cells at the 4- to 8-cell stage. A) Schematic depicting the canonical insulin/insulin-like growth factor signaling (IIS) pathway in *C. elegans*. **B)** Representative single plane images showing the localization of endogenously-tagged FoxO^DAF-16^::GFP (green in merged images) relative to mCherry::H2B (magenta in merged images) in control 2-cell through 32- to 64-cell stage embryos and *FoxO^daf-16^(RNAi)* 4-cell embryos; black box behind some images for display; scale bar=10 μm. **C)** Representative single plane images showing autofluorescence levels in a control N2 4-cell embryo using the same imaging settings in **(B)**; scale bar=10 μm. **D)** Graph plotting the average (avg.) nuclear levels of FoxO^DAF-16^::GFP in *FoxO^daf-16^(RNAi)* and control early worm embryos normalized (norm.) to the nuclear autofluorescence levels in N2 4-cell embryos. Error bars=SD; N=number of experimental replicates; n=number of nuclei scored for each genotype, developmental stage, and relative nuclear FoxO^DAF-16^::GFP levels (low or high) by color; n.s.=p-value not significant (>0.05), **=p-value ≤0.01, ***=p-value ≤0.001, and ****=p-value ≤0.0001 (1-way ANOVA; see **Table S1**).

While the role of FoxO^DAF-16^ in larval and adults is well characterized, how FoxO^DAF-16^ functions during early embryogenesis remains unknown. In mice, members of the FoxO family are necessary for the proper preimplantation embryonic development (Hosaka et al., 2004; Kuscu et al., 2019). In *Drosophila*, upregulation of FoxO decreases embryo size (Shrivastava et al., 2023). In *C. elegans*, FoxO^DAF-16^ regulates reproductive healthspan (Qin and Hubbard, 2015). However, the localization and/or role for FoxO^DAF-16^ in early worm embryos is not known.

Here we studied the inheritance pattern and role for FoxO^DAF-16^ in cleavage stage *C. elegans* embryos. Using quantitative live cell imaging of endogenously-tagged FoxO^DAF-16^::GFP, we show a previously uncharacterized expression pattern of FoxO^DAF-16^ during early embryogenesis. In 2- to 4-cell stage embryos, FoxO^DAF-16^ was present at uniformly low levels in somatic and germ precursor cell nuclei. At the 8-cell stage, FoxO^DAF-16^ became specifically enriched in germ precursor cell nuclei (P lineage) and was lost from somatic daughter cell progeny. This nuclear enrichment of FoxO^DAF-16^ in germ precursor cells was dependent on the master germ fate determinants PIE-1 and POS-1, the phospholipid regulators PI3K^AGE-1^ and PTEN^DAF-18^, and the deubiquitylase USP7/MATH-33, but surprisingly independent of the sole worm insulin receptor InR^DAF-2^ and AKT family kinases. Cell cycle timing analysis revealed a pace setter role for FoxO^DAF-16^ in slowing down cell cycle progression during early cleavage divisions, especially in the germ precursor cells. Together, our results describe the cell cycle role and early inheritance pattern of nuclear FoxO^DAF-16^, and suggests that some, but not all, canonical IIS pathway components contribute to this spatial patterning of nuclear FoxO^DAF-16^ during early embryogenesis.

## Results and Discussion

### FoxO^DAF-16^ is enriched in germ precursor cell nuclei during early embryogenesis

To characterize FoxO^DAF-16^ expression and localization during early *C. elegans* development, we used spinning disk confocal microscopy to image endogenously-tagged FoxO^DAF-16^::GFP in oocytes and early embryos from the 2-cell to 64-cell stage. For this, we generated a strain co-expressing endogenously-tagged FoxO^DAF-16^::GFP (Aghayeva et al., 2020) and transgenic mCherry::HistoneH2B (mCh::H2B (Audhya et al., 2005)), to label the nuclei. To control for background autofluorescence, we performed the same analysis in ancestral (wild-type) N2 Bristol embryos, which do not express any fluorescently-tagged reporters. In oocytes, FoxO^DAF-16^ was enriched in −5+ oocyte nuclei but was no longer detectable after oocyte maturation in −1 to −4-oocyte nuclei (proximal to the spermatheca), as has been previously reported (Zhang et al., 2022). No nuclear FoxO^DAF-16^ was detected in germline cells or oocytes after *FoxO^DAF-16^(RNAi)* (**Figure S1**; for review on the syncytial worm germline see (Hubbard, 2007; Hubbard and Greenstein, 2005; Pazdernik and Schedl, 2013)). In 2-cell and 4-cell stage embryos, we observed low levels of nuclear enrichment in all cells in FoxO^DAF-16^::GFP expressing embryos, but not after *FoxO^DAF-16^(RNAi)* or in wild-type N2 embryos (**Figure 1B-D**, **S2A-B**). Starting in 8-cell stage embryos, we no longer observed nuclear FoxO^DAF-16^ in cells in the embryo anterior but observed increased nuclear levels in only 1 or 2 cells in the embryo posterior (**Figure 1B, D**). Between the 16-cell and 64-cell stages, nuclear FoxO^DAF-16^ levels remained high in 1 or 2 posterior cells and increased as cell size decreased at each cleavage stage (**Figure 1B, D**). These results suggest that FoxO^DAF-16^ is preferentially inherited by a single posterior cell lineage and localizes to the nucleus during early worm embryogenesis.

To identify the posterior cells with high nuclear FoxO^DAF-16^ levels, we performed time-lapse cell lineage tracking. The early *C. elegans* cell lineage map is well characterized and invariant from embryo to embryo, and each cell is easily identifiable (Sulston et al., 1983) (**Figure 2A**). We used transmitted light (DIC) and mCh::H2B to track each cell lineage from 4-cell stage through 32-cell stage embryos, and imaged FoxO^DAF-16^ after each round of cleavage divisions. Relative to anterior somatic cells (ABal/r (8- to-16-cell) and ABal/ra/p (16- to 32-cell), hereafter ABax and ABaxx, respectively), nuclear FoxO^DAF-16^ levels were ∼1.5 fold higher in the P3 and C cells (8- to-16-cell stage) and ∼3 fold higher in the P4 and D cells (16- to 32-cell stage) (**Figure 2B-E**), confirming our initial observation that nuclear FoxO^DAF-16^ levels increase at each cleavage stage (**Figure 1B, D**). The P3 and P4 cells are cells in the P lineage or germ lineage, and their somatic sister cells are C and D, respectively (**Figure 2A**). No nuclear FoxO^DAF-16^ signal was observed in the daughters of C after it divided (**Figure 2B-E**), suggesting that FoxO^DAF-16^ is degraded (or excluded from the nuclei) in this somatic cell around mitosis (see more discussion below). Thus, we found that nuclear FoxO^DAF-16^ enrichment is inherited through the P lineage, which forms all gametes in the adult worm (Sulston et al., 1983).

**Figure 2:**
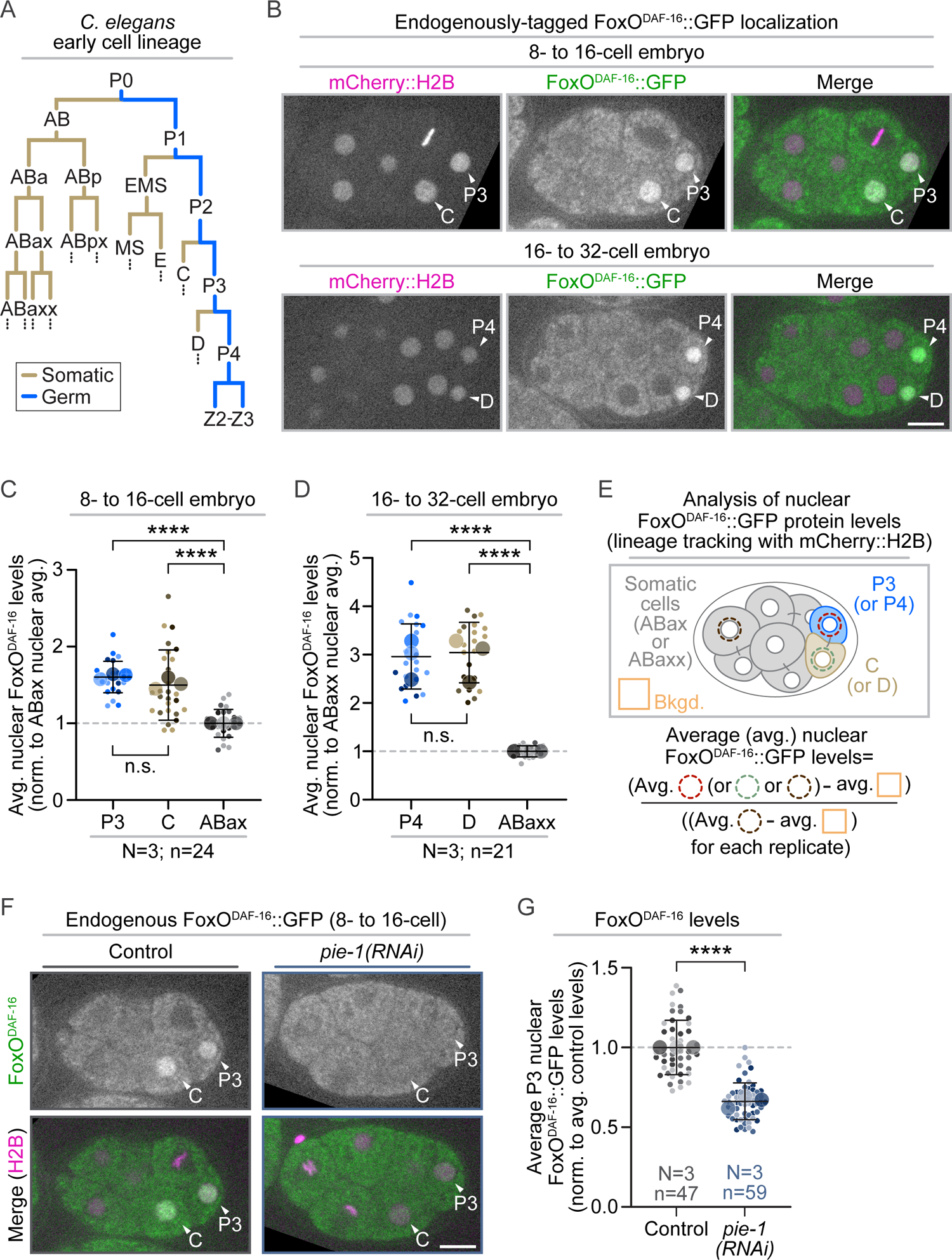
FoxODAF^-16^ is enriched in the nucleus of early germ precursor cells via a germ fate dependent mechanism. **A)** Schematic depicting the early *C. elegans* somatic (tan) and germ (blue) cell lineages. **B)** Representative single plane images after lineage tracking (from the 4-cell stage) using mCherry::H2B (magenta in merged images) showing nuclear enrichment of endogenously-tagged FoxO^DAF-16^::GFP in the P3 and C cells (8- to 16-cell embryos, top panels) and P4 and D cells (16- to 32-cell stage embryos, bottom panels); black box behind some images for display; scale bar=10 μm. **C)** Graph plotting the average (avg.) nuclear levels of FoxO^DAF-16^::GFP in the P3 (blues), C (browns), and other somatic ABax cells (grays) in 8- to 16-cell stage embryos normalized (norm.) to the average nuclear levels in somatic cells. **D)** Graph plotting the average nuclear levels of FoxO^DAF-16^::GFP in the P4 (blues), D (browns), and other somatic ABaxx cells (grays) in 16- to 32-cell stage embryos normalized to the average nuclear levels in somatic cells. (**C-D**) Small circles indicate the average levels in individual nuclei and large circles and indicate replicate averages; shading indicates results from the same replicate (*e.g.*, lightest shades in P3, C and somatic ABax cell conditions from same replicate). Error bars=SD; N=number of experimental replicates; n=number of nuclei scored for each cell type by color; n.s.=p-value not significant (>0.05), ****=p-value ≤0.0001 (1-way ANOVA; see **Table S1**). **E)** Schematic depicting analysis shown in **(C-D)** performed on single plane images to quantify nuclear FoxO^DAF-16^::GFP levels in the germ precursor cells (P3 and P4, dashed red circle), their somatic sister cells (C and D, dashed green circle), other somatic cells (dashed brown circle), and average extracellular background signal (orange box); see also Methods. **F)** Representative single plane images showing FoxO^DAF-16^::GFP (green in merged images) and mCherry::H2B (magenta in merged images) localization in 8- to 16-cell embryos with and without *pie-1(RNAi)*; black box behind some images for display; scale bar=10 μm. **G)** Graph plotting the average P3 nuclear levels of FoxO^DAF-16^::GFP in control (grays) and *pie-1(RNAi)* (blues) embryos. Results are normalized to the average nuclear levels in control embryos. Small circles indicate the average levels in individual P3 nuclei and large circles indicate replicate averages; shading indicates results from same replicate (*e.g.*, lightest shades in both control and RNAi conditions from the same replicate). Error bars=SD; N=number of experimental replicates; n=number of P3 nuclei scored for each genotype by color; ****=p-value ≤0.0001 (Student’s t-test, unpaired; see **Table S1**).

### Germ fate determinants are required for nuclear FoxO^DAF-16^ in the P3 germ precursor cell

We next tested if germ fate specification is required for nuclear enrichment of FoxO^DAF-16^ in germ precursor cells. The CCCH Zn-finger protein PIE-1 (Pharynx and Intestine in Excess) is the master regulator of germ fate determination in *C. elegans*. (Bowerman, 1995; Mango et al., 1994; Mello et al., 1992; Mello et al., 1996; Strome, 2005) POS-1 (Posterior Segregation), a homolog of PIE-1, is also required for germ fate specification and important for proper PIE-1 localization (Tabara et al., 1999). In the early embryo, both PIE-1 and POS-1 are asymmetrically inherited specifically by germ precursor cells (P lineage), where they regulate a variety of germ-cell specific processes including transcription, gene silencing, and translation. (Batchelder et al., 1999; Bowerman, 1995; D’Agostino et al., 2006; Ghosh and Seydoux, 2008; Kim et al., 2021; Mango et al., 1994; Mello et al., 1992; Mello et al., 1996; Ogura et al., 2003; Reese et al., 2000; Seydoux, 1996; Seydoux and Dunn, 1997; Tenenhaus et al., 1998; Tenenhaus et al., 2001; Wu et al., 2015; Zhang et al., 2003) To determine if germ fate is required for nuclear FoxO^DAF-16^ enrichment in germ precursor cells, we measured nuclear levels of endogenously-tagged FoxO^DAF-16^::GFP in the P3 cell with and without RNAi-mediated depletion of PIE-1 and POS-1. Relative to control embryos, P3 nuclear FoxO^DAF-16^ levels were reduced in both *pie-1(RNAi)* embryos (30-38% lower in *pie-1(RNAi)* replicates than in controls, **Figure 2F-G**) and *pos-1(RNAi)* embryos (15-20% lower in *pos-1(RNAi)* replicates than in controls, **Figure S2C-D**). Thus, proper germ cell fate specification is required for nuclear enrichment of FoxO^DAF-16^ (or protein stability) in the P3 germ precursor cell.

Given that PIE-1 is important for enriched FoxO^DAF-16^ nuclear levels in the germ lineage, we also tested if FoxO^DAF-16^ is inversely important for PIE-1 levels and localization. Unlike PIE-1, FoxO^DAF-16^ is not essential for germline fate specification (*e.g.*, see (Qin and Hubbard, 2015)), but has been reported to regulate PIE-1 expression in some contexts ((Curran et al., 2009), in contrast see (Knutson et al., 2016)). To test if FoxO^DAF-16^ regulates PIE-1 expression in early embryos, we depleted FoxO^DAF-16^ using RNAi in a strain expressing endogenously-tagged PIE-1::GFP (Kim et al., 2014) and quantified PIE-1 levels in the P2, P3, and P4 germ precursor cells (4-cell through 32-cell stage embryos). We did not observe any significant difference between PIE-1 levels (or localization) at any stage in control or *FoxO^daf-16^(RNAi)* embryos (**Figure S3**). This suggests that FoxO^DAF-16^ does not regulate PIE-1 levels in the early worm embryo.

### Nuclear FoxO^DAF-16^ enrichment in the P3 germ precursor cell is controlled by phospholipid regulators

We next tested if the observed inheritance pattern for FoxO^DAF-16^ during embryogenesis is regulated by the canonical worm insulin/insulin-like growth factor signaling (IIS) pathway (**Figure 1A**). We took a systematic approach by depleting all IIS pathway components individually or in combination using RNAi and quantifying the average nuclear levels of endogenously-tagged FoxO^DAF-16^::GFP in the P3 cell (8- to 16-cell stage embryos) with and without RNAi knockdown. For each experimental replicate, a similar number of control and IIS pathway component RNAi embryos were collected to allow individual replicate (**Figure S4**) and pooled replicate comparisons (**Figure 3**). RNAi depletion of PI3K^AGE-1^, the lipid kinase that phosphorylates PIP_2_ to make PIP_3_, significantly reduced P3 nuclear levels of FoxO^DAF-16^ (20-25% lower in *PI3K^AGE-1^(RNAi)* replicates than in controls, **Figure 3A-B, S4**). Depletion of PTEN^DAF-18^, the lipid phosphatase that converts PIP_3_ back to PIP_2_, also decreased P3 nuclear FoxO^DAF-16^ levels (16-25% lower in *PTEN^DAF-18^(RNAi)* replicates than in controls, **Figure 3A-B, S4**). Depletion of PDK-1, which is normally activated by PIP_3_, led to a partial reduction in P3 nuclear FoxO^DAF-16^ levels (∼8% lower in *pdk-1(RNAi)* versus in replicate controls, **Figure S4**), but the difference was not significant when compared to pooled controls (**Figure 3A, C**). Worms have three AKT homologs: AKT-1, AKT-2, and SGK-1 (**Figure 1A**); while *akt-1* is expressed at high levels in early embryos, *akt-2* is only detectable in a few anterior cells, and *sgk-1* is not detectably expressed (Tintori et al., 2016). There was no difference in P3 nuclear FoxO^DAF-16^ levels with or without RNAi-mediated depletion of AKT-1 either alone or in combination with AKT-2 and/or SGK-1 (**Figure 3A, C, S4**). There was also no difference in P3 nuclear FoxO^DAF-16^ levels with or without *InR^daf-2^(RNAi)* (**Figure 3A-B, S4**), suggesting this early FoxO^DAF-16^ patterning is also insulin receptor-independent. Taken together, this data suggests that early embryonic FoxO^DAF-16^ is only regulated by some members of the IIS pathway and that phospholipid regulation is primarily responsible for maintaining nuclear FoxO^DAF-16^ levels in the germ lineage.

**Figure 3:**
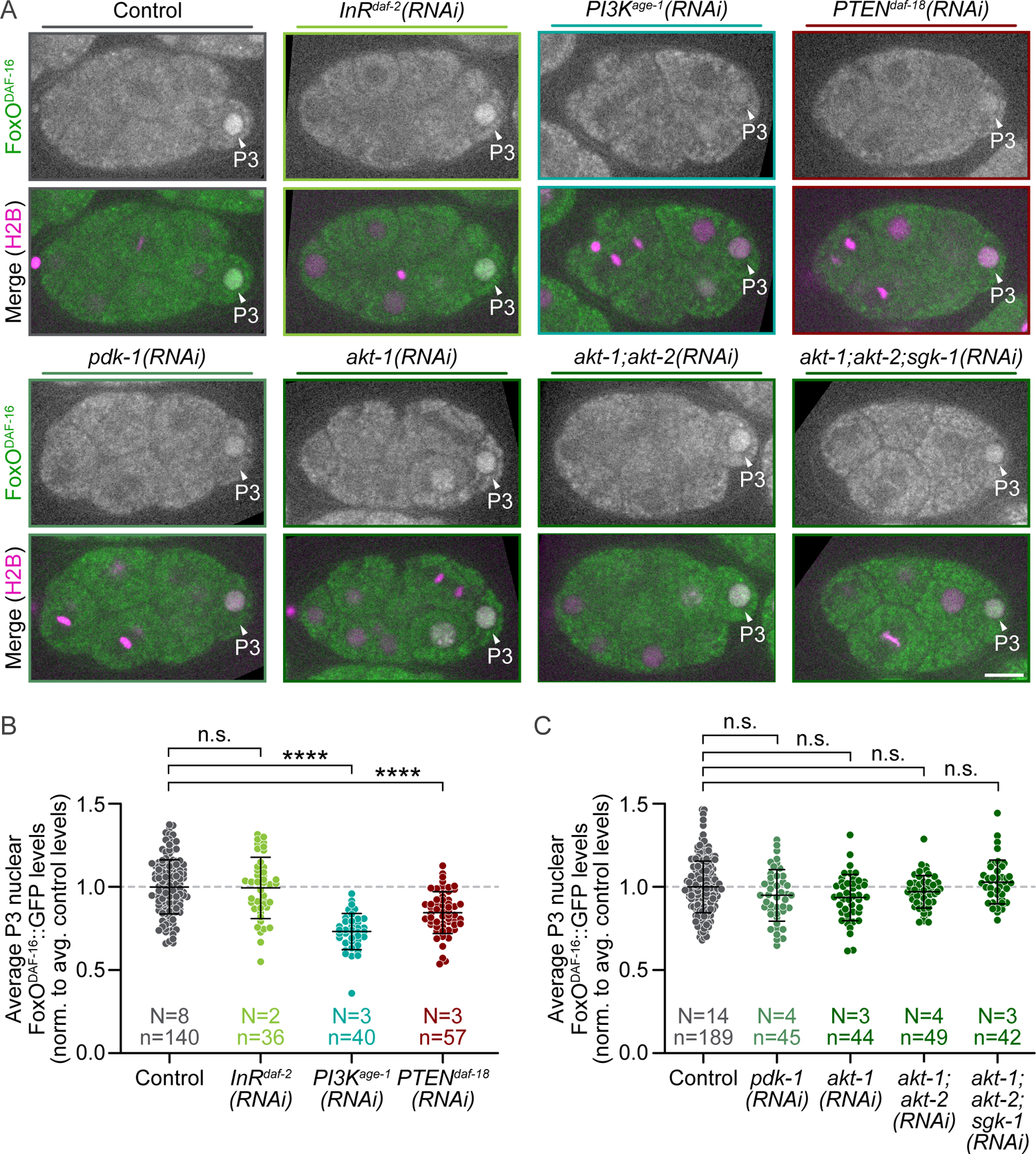
PI3KAGE^-1^ and PTEN^DAF-18^ are required for robust FoxO^DAF-16^ nuclear enrichment in the P3 germ precursor cell. A) Representative single plane images showing FoxO^DAF-16^::GFP (green in merged images) and mCherry::H2B (magenta in merged images) localization in 8- to 16-cell embryos with and without RNAi-mediated knockdown of insulin/insulin-like growth factor signaling (IIS) pathway components InR^DAF-2^, PI3K^AGE-1^, PTEN^DAF-18^, PDK-1, and AKT family homologs (AKT-1, AKT-2, and SGK-1) alone and together (for schematic of IIS pathway see Figure 1A); black box behind some images for display; scale bar=10 μm. B-C) Graphs plotting the average P3 nuclear levels of endogenously tagged FoxO^DAF-16^::GFP with and without RNAi-mediated knockdown of IIS pathway components; results from each genotype are normalized (norm.) to the average (avg.) nuclear levels in pooled controls for each graph (for comparisons with individual controls for each replicate, see Figure S4). Error bars=SD; N=number of experimental replicates; n=number of P3 nuclei scored for each genotype by color; n.s.=p-value not significant (>0.05), *=p-value ≤0.05, **=p-value ≤0.01, and ****=p-value ≤0.0001 (1-way ANOVA; see Table S1).

Our observation that depletion of AKT family kinases by RNAi had no effect on FoxO^DAF-16^ was initially puzzling. However, recent work in human breast cancer cells showed AKT only regulates FoxO in a stimulus-dependent manner (Lasick et al., 2023). Specifically, AKT-based phosphorylation only impacts nuclear localization of FoxO under starvation conditions, but not when cells are treated with H_2_O_2_ to stimulate oxidative stress (Lasick et al., 2023). AKT-independent regulation of nuclear FoxO has also been widely found in hematopoietic stem and progenitor cells (Liang et al., 2016), human primary breast tumor cells (Hu et al., 2004), prostate cancer cells (Li et al., 2003), acute myeloid leukemia cells (Chapuis et al., 2010), glioblastoma cells (Lau et al., 2009; Masui et al., 2013), and human embryonic stem cells (Zhang et al., 2011). Thus, it seems that phosphorylation by AKT family kinases is not the only method of regulating FoxO nuclear levels, especially in cancer cells and in cell types with high potency, such as stem and progenitor cells, including *C. elegans* germ precursor cells.

In *C. elegans*, AKT-independent FoxO^DAF-16^ regulation is mediated by several signaling pathways (Sun et al., 2017; Tissenbaum, 2018). For example, JNK-1 (c-Jun N-terminal kinase) directly interacts with and phosphorylates FoxO^DAF-16^ to regulate lifespan and in response to heat stress (Oh et al., 2005). FoxO^DAF-16^ is also regulated by AMP kinase (Greer et al., 2007) and the TOR pathway (target of rapamycin) in response to environmental ques (Robida-Stubbs et al., 2012). To determine if either the JNK-1 or TOR pathways regulate nuclear enrichment of FoxO^DAF-16^ in the germ lineage, we depleted *jnk-1* and the key rapamycin-binding TORC1 (TOR complex 1) component, *Raptor^daf-15^*, using RNAi and measured the average nuclear levels of endogenously-tagged FoxO^DAF-16^::GFP in the P3 cell. We observed no difference in P3 nuclear FoxO^DAF-16^ levels between control and *jnk-1* or *Raptor^daf-15^* depleted embryos (**Figure S5A-D**). Thus, FoxO^DAF-16^ nuclear enrichment in the germ lineage is also independent of JNK-1 and TORC1.

It is unclear what activates PI3K^AGE-1^ and/or PTEN^DAF-18^ to control FoxO^DAF-16^ levels in the early germ precursor cells. One candidate tyrosine kinase is MES-1, a protein that like PIE-1, is important for germ fate specification in the early worm embryo (Bei et al., 2002; Berkowitz and Strome, 2000). However, because of MES-1’s role in germ fate specification, teasing apart germ fate-mediated FoxO^DAF-16^ regulation from a direct role for MES-1 in phospholipid-mediated FoxO^DAF-16^ regulation within the germ precursor cells will be challenging. In future research, the *C. elegans* early embryo may serve as a system to genetically screen for AKT-independent regulators of FoxO^DAF-16^ nuclear enrichment.

### Nuclear FoxO^DAF-16^ enrichment in the P3 germ precursor cell is controlled by the deubiquitinylase USP7/MATH-33

We next tested if ubiquitinylation regulates FoxO^DAF-16^ P3 nuclear levels. In several models, proteasome inhibition leads to stabilization of FoxO levels, suggesting it is regulated by active degradation. (Aoki et al., 2004; Hu et al., 2004; Matsuzaki et al., 2003; Plas and Thompson, 2003) In *C. elegans* aging, FoxO^DAF-16^ is ubiquitinylated and targeted for degradation by the RC3H1-like E3 ubiquitin ligase RLE-1 (Li et al., 2007). This is countered by the *C. elegans* homolog of the USP7-like deubiquitinylase, MATH-33 (Heimbucher et al., 2015). To test the role of these ubiquitin regulators in FoxO^DAF-16^ patterning during early embryogenesis, we used RNAi to deplete RLE-1 and MATH-33 and measured the levels of endogenously tagged FoxO^DAF-16^::GFP in P3 cell nuclei. RNAi depletion of *rle-1* did not alter P3 nuclear FoxO^DAF-16^ levels (**Figure S5E-F**). In contrast, RNAi depletion of *math-33* dramatically lowered P3 nuclear FoxO^DAF-16^ levels (∼28-40% lower in *math-33(RNAi)* replicates than in controls, **Figure 4A-B**). These data suggest that the deubiquitinylase MATH-33 regulates FoxO^DAF-16^ during early embryogenesis. The specific E3 ligase(s) that counteracts MATH-33 activity at this stage remains unknown.

**Figure 4:**
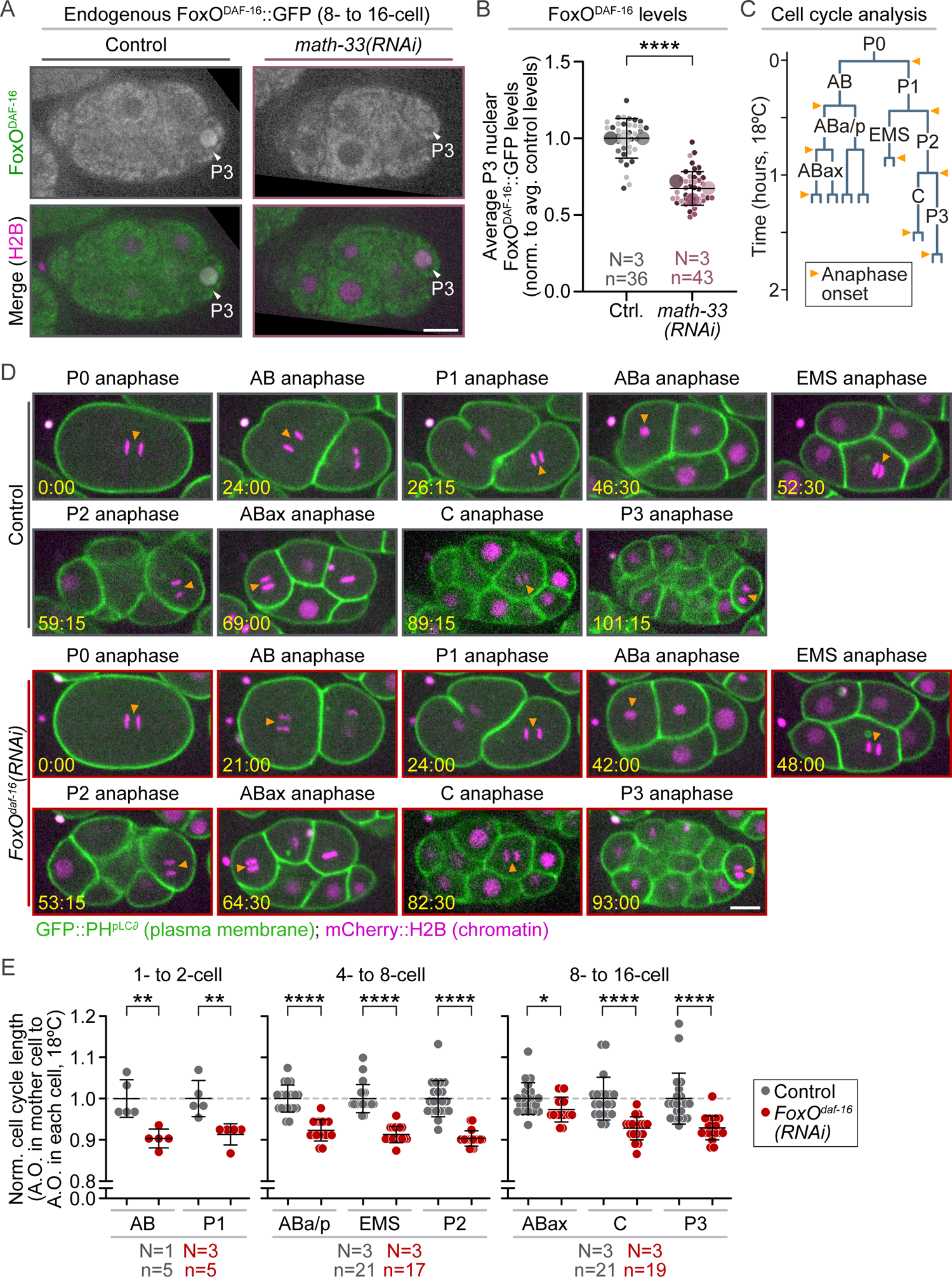
FoxODAF^-16^ in P3 is dependent on the deubiquitylase MATH-33 and slows cell cycle timing in the early embryo. **A)** Representative single plane images showing FoxO^DAF-16^::GFP (green in merged images) and mCherry::H2B (magenta in merged images) localization in 8- to 16-cell embryos with and without *math-33(RNAi)*; black box behind some images for display; scale bar=10 μm. **B)** Graph plotting the average P3 nuclear levels of FoxO^DAF-16^::GFP in control (grays) and *math-33(RNAi)* (purples) embryos. Results are normalized (norm.) to the average (avg.) nuclear levels in control embryos. Small circles indicate the average levels in individual P3 nuclei and large circles indicate replicate averages; shading indicates results from same replicate (*e.g.*, lightest shades in both control and RNAi conditions from the same replicate). Error bars=SD; N=number of experimental replicates; n=number of P3 nuclei scored for each genotype by color; ****=p-value ≤0.0001 (Student’s t-test, unpaired; see **Table S1**). **C)** Schematic of the early *C. elegans* cell lineage (drawn to scale with time from anaphase onset in P0) indicating the time points scored for anaphase onset in each cell (orange arrowheads). **D)** Representative single plane images of embryos expressing GFP::PH^pLCδ^ (plasma membrane, green in merged images) and mCherry::H2B (magenta in merged images) showing the timing of anaphase onset in each indicated cell during early worm development at 18°C; orange arrowheads indicate anaphase onset in that cell; scale bar=10 μm; time (minutes:seconds post-anaphase in P0) is in yellow. **E)** Graph plotting the average cell cycle timing at 18°C (cell cycle time=anaphase onset (A.O.) in mother cell to A.O. in each indicated cell) for individual cells in control (gray) and *FoxO^daf-16^(RNAi)* (red) embryos normalized to the average cell cycle timing in controls. Results are shown for the 2- to 4-cell embryo (AB and P1; far left graph), 4- to 8-cell (ABa/p, EMS, and P2; center graph), and 8- to 16-cell embryo (ABax, C, and P3; right graph). Error bars=SD; N=number of experimental replicates; n=number of embryos analyzed for each genotype by color; *=p-value ≤0.05, **=p-value ≤0.01, ****=p-value ≤0.0001 (Student’s t-test, unpaired; see **Table S1**).

A strong candidate E3 ubiquitin ligase for regulating FoxO^DAF-16^ levels in the somatic sisters of germ precursor cells is the <u>A</u>naphase <u>P</u>romoting <u>C</u>omplex/<u>C</u>yclosome (APC/C), which is essential for proper cell-cycle progression (Peters, 2006). Supporting this hypothesis, recent work in human urothelial bladder cancer found that the APC/C ubiquitinylates FoxO3, leading to its cell cycle-dependent degradation (Yan et al., 2023). In our lineage studies, we found that after germ precursor cell division at the 8- to 16-cell stage, nuclear FoxO^DAF-16^ levels were initially equal in both the somatic C and germ lineage P3 daughter cells (**Figure 2**). However, after cell division in the somatic C cell nuclear FoxO^DAF-16^ levels did not return, suggesting it is degraded at some point near or during M-phase in C. The APC/C promotes anaphase onset and exit from M-phase (Peters, 2006); thus, peak APC/C activity is temporally coordinated with FoxO^DAF-16^ degradation in these cells. Better tools to allow acute regulation of the APC/C (and early cell cycle progression) will help to determine how ubiquitinylation and/or other post-translational modifications regulate FoxO^DAF-16^ levels in the early embryo.

### FoxO^DAF-16^ slows down cell cycle progression during early embryogenesis

One of the first identified functions for FoxO was in cell cycle regulation during G1/S and G2/M transitions. (Alvarez et al., 2001; Dijkers et al., 2000; Medema et al., 2000) During larval *C. elegans* development, FoxO^DAF-16^ is required for starvation-dependent cell cycle arrest of germline progenitor cells Z2-Z3, the daughters of the P4 cell. (Baugh and Sternberg, 2006) In adults, FoxO^DAF-16^ regulates cell cycle arrest to prevent oogenesis in the germline upon DNA damage. (Sarkar et al., 2023) FoxO also plays a role in cell cycle progression and quiescence in hydra stem cells (Boehm et al., 2012), the *Drosophila* stem cell niche (Yang et al., 2013), mouse adult muscle stem cells (Gopinath et al., 2014), and human embryonic stem cells (Zhang et al., 2011). Thus, we reasoned that FoxO^DAF-16^ may also regulate cell cycle progression during early worm embryogenesis.

To test if FoxO^DAF-16^ regulates cell cycle progression in early worm embryos, we used time-lapse spinning disc confocal microscopy to monitor the timing of each cell division in embryos co-expressing GFP::PH^pLCδ^ to label the plasma membrane and mCh::H2B to label the chromatin (Audhya et al., 2005) with and without *FoxO^daf-16^(RNAi)*. We tracked the time between anaphase onset in each cell from the 1-cell through the 8- to 16-cell stage at 18°C with 45-second temporal resolution (**Figure 4C**). At the 2- to 4-cell and 4- to 8-cell stage, all cells in *FoxO^daf-16^(RNAi)* embryos progressed through the cell cycle ∼7-10% more quickly than in control embryos (**Figure 4D-E, S6**). At the 8- to 16-cell stage, all cells also progressed through the cell cycle more quickly in *FoxO^daf-16^(RNAi)* embryos. Unlike earlier stages, at this stage of development, when FoxO^DAF-16^ becomes enriched in the P3 and C cells, we found that this cell cycle effect was cell type-specific: the cell cycle was only ∼3% faster in the anterior somatic ABax cells, whereas both the P3 germ precursor and its somatic sister C both divided ∼7% faster in *FoxO^daf-16^(RNAi)* embryos than in controls (**Figure 4D-E, S6**). Thus, in early worm embryos, FoxO^DAF-16^ functions as a cell cycle pacer to slow down the early cleavage divisions.

Taken together, our results show that FoxO^DAF-x16^ is expressed and asymmetrically inherited in the germ precursor cells via a phospholipid regulator and germ fate-dependent process. We also found that FoxO^DAF-16^ functions as a cell cycle pacer to slow cell cycle timing during the early worm cleavage divisions. Future studies will further elucidate the cell and molecular mechanisms that underly asymmetric FoxO^DAF-16^ inheritance specifically in the germ lineage. It will also be interesting to determine if, like in adult worms (Leiser et al., 2011; Sarkar et al., 2023), nuclear FoxO^DAF-16^ functions to protect germ precursor cells specifically and/or other cells in the early embryo from environmental stressors (*e.g.*, DNA damaging agents or high temperatures).

## Materials and Methods

Much of these materials and methods are adapted from (Connors et al., 2024). Wormbase (wormbase.org (Davis et al., 2022)) was used extensively in this work.

### Worm strain maintenance

The *C. elegans* strains used in this study are indicated in **Table S1** and were grown similar to as described (Brenner, 1974). Briefly, worm strains were grown on non-vented standard 60 mm plates (T3308, Tritech Research). Each plate was filled with 10.5 mL nematode growth media (NGM) (23 g Nematode Growth Medium (Legacy Biologicals, a division of Research Products International), 1 mL 1M CaCl_2_, 1 mL of 1M MgSO_4_, 25 mL of 1M K_3_PO_4,_ 975 mL ddH_2_O) using a PourBoy 4 (PB4, Tritech Research) and seeded with seeded with 500 μL OP50 *E. coli* bacteria as a food source. Strains were maintained at 20°C in cooling/heating incubators (Binder).

### RNA-mediated interference (RNAi) by injection

For all RNAi experiments in this manuscript, dsRNA was injected into L4 stages hermaphrodites 24 hours prior to imaging. For each target gene, ∼500-1000 bp of gene sequence was PCR amplified using primers containing a T7 promoter sequence and cDNA as a template (when possible, this region was within a single exon). Primers used to generate dsRNA for each target gene are listed in **Table S1**; E-RNAi (Horn and Boutros, 2010) or Primer3 (Untergasser et al., 2012) were used to design primers. PCR products were confirmed with a 1% agarose gel and PCR purified (QIAquick PCR Purification kit, QIAGEN). The dsRNA was synthesized using a T7 reverse transcription reaction kit (MEGAscript, Life Technologies) and purified using phenol-chloroform. For phenol-chloroform purifications, the synthesized ssRNA was mixed 1:1 with phenol-chloroform (Invitrogen), vortexed for 2 minutes, and spun down for 3 minutes at 12,000 x g in a Beckman Coulter microfuge 16. The aqueous layer was then transferred to new tube, mixed again 1:1 with phenol-chloroform, vortexed for 2 minutes, and spun down for 3 minutes at 12,000 x g in a Beckman Coulter microfuge 16. The aqueous layer was transferred to a new tube, mixed 1:1 with pre-chilled (−20°C) isopropanol (100%, Sigma), and incubated at −20°C overnight (∼16 hours). The ssRNA was precipitated by spinning the tube down at 12,000 x g for 15 minutes in a Beckman Coulter microfuge 16. The pellet (translucent/clear/white) was allowed to air dry for ∼5 minutes and then resuspended in 1x soaking buffer (32.7 mM Na_2_HPO_4_, 16.5 mM KH_2_PO_4_, 6.3 mM NaCl, 14.2 mM NH_4_Cl). To make dsRNA, ssRNA reactions were combined and annealed at 68°C for 10 minutes followed by 37°C for 30 minutes. The dsRNAs were diluted to a final concentration of ∼2000-2500 ng/μL (when possible) and 2 μL aliquots of the dsRNA were stored at −80°C until needed.

For each experiment, a fresh aliquot (or aliquots for double or triple RNAi experiments) was diluted to ∼1000 ng/μL (∼1000 ng/μL for *akt-1* and ∼500 ng/μL for *akt-2* dsRNAs in double RNAi experiments; ∼750 ng/μL for *akt-1*, ∼375 ng/μL for *akt-2*, and ∼375 ng/μL for *sgk-1* dsRNAs in triple RNAi experiments) using 1x soaking buffer and centrifuged at 16,000 x g for 10 minutes at room temperature (∼22°C) in a Beckman Coulter microfuge 16 prior to use. Machine pulled borosilicate glass capillary needles were made fresh every week (World Precision Instruments, WPI; Sutter Instruments, P1000 needle puller). 0.4 μL of the diluted dsRNA was loaded into the back of pulled borosilicate glass capillary needles and injected into the gut of L4 hermaphrodites using a Leica DMIRB microscope equipped with Hoffman optics, a Plan L 20x/0.4 CORR PH (Leica), a rotating stage, and the XenoWorks digital microinjector and micromanipulator injection system (Sutter Instruments). Groups of 5-10 worms were injected at a time, rescued by resuspension in M9 buffer (6 g KH_2_PO_4_, 12 g Na_2_HPO_4_, 10 g NaCl, 0.5 mL 1 M MgSO_4_, ddH_2_O to 2 L) to fresh plates seeded with OP50 bacteria, and allowed to recover for ∼24 hours at 20°C prior to imaging.

### Worm and embryo preparation for live cell imaging

Young gravid adult hermaphrodites were dissected on a high-resolution dissecting microscope (Olympus SZX16 with an Olympus SDF PLAPO 1XPF objective) in M9 buffer. Embryos were mounted on a thin (∼1-2x lab tape thickness) 2% agar pad on a glass slide (VWR VistaVision, 3 inches x 1 inch x 1 mm) using a hand pulled borosilicate glass capillary pipette (World Precision Instruments, WPI) as a mouth pipette. To image the germline and oocytes in adults (**Figure S1**), worms were first anesthetized in 1x levamisole (0.01%) prior to mounting on the agar pad. A 22 x 22 mm No. 1.5 glass coverslip (VWR) was placed on top of the embryos or worms (germlines/oocytes) for imaging, similar to as described (Gonczy et al., 1999).

### Live cell imaging microscope set up and temperature control

For all imaging experiments, room and microscope temperatures were continuously monitored using a Bluetooth-enabled smart temperature sensor (SensorPush) on the microscope stage. All imaging was done at 26 ± 0.5°C except for the cell cycle timing analysis (**Figure 4D-E, S6**), which was done at 18 ± 0.5°C.

For all experiments except in **Figure S3A**, we used an inverted Nikon Eclipse Ti microscope with a spinning disc confocal unit (Yokogawa, CSU-10 with Borealis upgrade (Spectral Applied Research)), a charge-coupled device (CCD) Orca-R2 camera (Hamamatsu Photonics), and a Piezo-driven (Applied Scientific Instrumentation, ASI) motorized stage for Z-sectioning. Focus was established and maintained using Nikon’s “Perfect Focus” before each Z-series was acquired. Two (488 nm and 561 nm) 150 mW Excitation lasers (ILE-2, Spectral Applied Research) were used and controlled by an acousto-optic tunable filter (Spectral Applied Research). A filter wheel (Sutter Instruments) was used for emission filter (525/50 nm and 620/60 nm bandpass (Chroma)) and DIC analyzer selection. All microscope systems and experimental setups were controlled and set up using the MetaMorph software (Molecular Devices). Room temperature was established and maintained using a heat pump-based temperature control device (MHWX, MultiAqua).

For experiments in **Figure S3A**, images were acquired on an inverted Nikon Eclipse Ti microscope (modified for compatibility with near-infrared light, as in (Hirsch et al., 2018; Sundaramoorthy et al., 2017)). The microscope was equipped with a spinning disc confocal unit (Yokogawa, CSU-10 with Borealis upgrade (Spectral Applied Research)), an Orca-R2 CCD camera (Hamamatsu Photonics), and a Piezo-driven (ASI) motorized stage for Z-sectioning. Two solid state (Cairn) 150 mW 488 nm and 561 nm lasers were used for excitation light, with a filter wheel (Ludl Instruments) that was used for DIC polarizer and emission filter (525/50 nm and 620/50 nm bandpass (Chroma)) selection. Room temperature was established and maintained using a heat pump-based temperature control (Mr. Slim, MSZ-D36NA; Mitsubishi).

### Live cell imaging and analysis parameters

All data analysis was performed using FIJI (FIJI is Just ImageJ) software (Schindelin et al., 2012).

### Quantitative analysis of endogenous FoxO^DAF-16^::GFP average nuclear levels

For quantitative imaging of average nuclear FoxO^DAF-16^ levels, we generated a strain expressing endogenously-tagged FoxO^DAF-16^::GFP (Aghayeva et al., 2020) and transgenic mCherry::H2B (mCh::H2B (Audhya et al., 2005)). Embryos were monitored every 60 seconds using mCh::H2B from the 4-cell stage until the 8-cell (**Figure 2-4A; S2C,** and **S5**) or the 16-cell (**Figure 2B**) stage, except in **Figure 1**, in which random groups of embryos at different stages were collected and imaged. At each time point we acquired an image using a 60x Plan Apo 1.40 N.A. oil immersion objective (Nikon) with 2 x 2 binning and 26 x 1.0 µm Z-sectioning of mCh::H2B to determine the cell cycle and developmental stage. Once the desired stage was reached (*e.g.*, after cell division of the P2 or P3 cell), we then acquired a single time point image stack as described above for both mCh::H2B and FoxO^DAF-16^::GFP.

For image analysis in **Figure 1D, 2C-D, G, 3B-C, 4B, S1B, S2B, D, S4,** and **S5B, D, F** the Z plane containing the most “in focus” image was selected and the nucleus was traced using the circle tool in FIJI. Next, a 50×50 pixel box was drawn outside of the embryo to measure the average extracellular background (camera background). This value was then multiplied by the area of the nucleus and subtracted from the total fluorescence intensity in the nucleus. This background subtracted value was then divided by the area of the nucleus to calculate the average nuclear fluorescence intensity. That data was then normalized by dividing each data-point by its replicate control average or pooled control average (except for **Figure 1D**, which was normalized to N2 levels) and plotted as superplots (Lord et al., 2020) except for **Figure 1D, 4E, S2B,** and **S6** in which only individual data points are shown. See also schematics in **Figure 2E, S1C** for a visual depiction of how quantitative image analysis of nuclear FoxO^DAF-16^::GFP levels was performed.

### Quantitative analysis of PIE-1::GFP P2, P3, and P4 cellular levels

For quantitative imaging of cellular levels of PIE-1 in P2, P3, and P4 cells we used a strain expressing endogenously-tagged PIE-1::GFP (Kim et al., 2021) (**Figure S3**). At each embryonic stage we captured an image using a 60x Plan Apo 1.40 N.A. oil immersion objective (Nikon) with 2 x 2 binning and 26 x 1.0 µm Z-sectioning. For image analysis, a sum projection of the embryo was generated and the entire P2, P3, or P4 cell was traced using the freehand tool in FIJI. To subtract background, a 20 x 20-pixel box was drawn in the cytoplasm of a non-enriched cell (somatic cell in the embryo anterior, ABa, ABax, or ABaxx). The data was then normalized by dividing by the control average for each replicate. See also schematic in **Figure S3C** for a visual depiction of how quantitative image analysis of PIE-1::GFP levels in germ precursor cells was performed

### Cell cycle timing analysis

To determine cell cycle timing (**Figure 4C-F, S6**), we used a strain expressing a marker for the plasma membrane (GFP::PH^pLCδ^) and for histone (mCh::H2B). Embryos were imaged from the 1-cell stage until ∼32-cell stage at 18 ± 0.5°C. Images were acquired every 45 seconds with a 40x Plan Fluor 1.30 N.A. oil immersion objective (Nikon), 2 x 2 binning, and 15 x 2 µm Z-sectioning. Cell cycle timing for each cell was measured from the time of anaphase onset (A.O.) in the mother cell to the time of anaphase onset in each cell (time from cell birth to cell division=cell cycle length). The data was then normalized by dividing by the average cell cycle length in controls. See also schematic in **Figure 4C** for a visual depiction of cell cycle timing analysis.

## Figure preparation

Figures for this manuscript were all generated using Adobe Illustrator 2023 (Adobe) and all graphs were created in Prism (Graphpad).

## Statistical analysis

All statistical analysis was performed using Prism 9 (Graphpad) or Excel (Microsoft). All RNAi experiments were repeated 2-4 times (experimental replicates; the number of experimental replicates (N) and cells/embryos analyzed (n) is indicated on each graph in all figures). For Figures 1D**, 2C-D, 3B-C,** and **S2B**, a 1-way ANOVA was used. For all other figures, a Student’s t-test was performed (unpaired). Error bars in all graphs represent the standard deviation (SD). p-values: n.s.=p>0.05, *=p≤0.05, **=p≤0.01, ***=p≤0.001, and ****=p≤0.0001. See **Table S1** for a detailed list of all statistical tests and p-values.

## Acknowledgements

We thank all members of the Canman and Shirasu-Hiza labs for their support, feedback, and advice on this work. We thank Dr. Elena Lucchetta for generating strain JCC1092. We are grateful for Adriana Hernandez and Michelle (Mimi) Schmidt for making worm plates and other essential lab reagents. We thank Drs. Eunhee Choi and Rebecca Haeusler for helpful discussions and suggestions. We are grateful to Dr. Oliver Hobert’s lab, Dr. Craig Mello’s lab, and the *Caenorhabditis* Genetics Center (NIH P40OD010440) for providing worm strains. We thank Hyo Taek Kim for color inspiration (Colors of Star Wars series). This work was funded by: NIH R01GM117407 (JCC), R01GM130764 (JCC), European Research Council CoG ChromoSOMe N°819179 (JD), NIH R01AG045842 (MSH), and NIH R35GM127049 (MSH). The authors declare no competing financial interests.

## Author contributions

M.S. Mauro and J.C. Canman conceived of the project and designed all experiments. M.S. Mauro and S.L. Martin made the dsRNAs for RNAi experiments. M.S. Mauro performed and analyzed all experiments except for the PIE-1::GFP imaging, which was performed by S.L. Martin, and analyzed by M.S. Mauro. M.S. Mauro, M. Shirasu-Hiza, J. Dumont, and J.C. Canman made significant intellectual contributions and helped write (or edit) the manuscript. M.S. Mauro and J.C. Canman made the figures.

---

**Figure S1:**
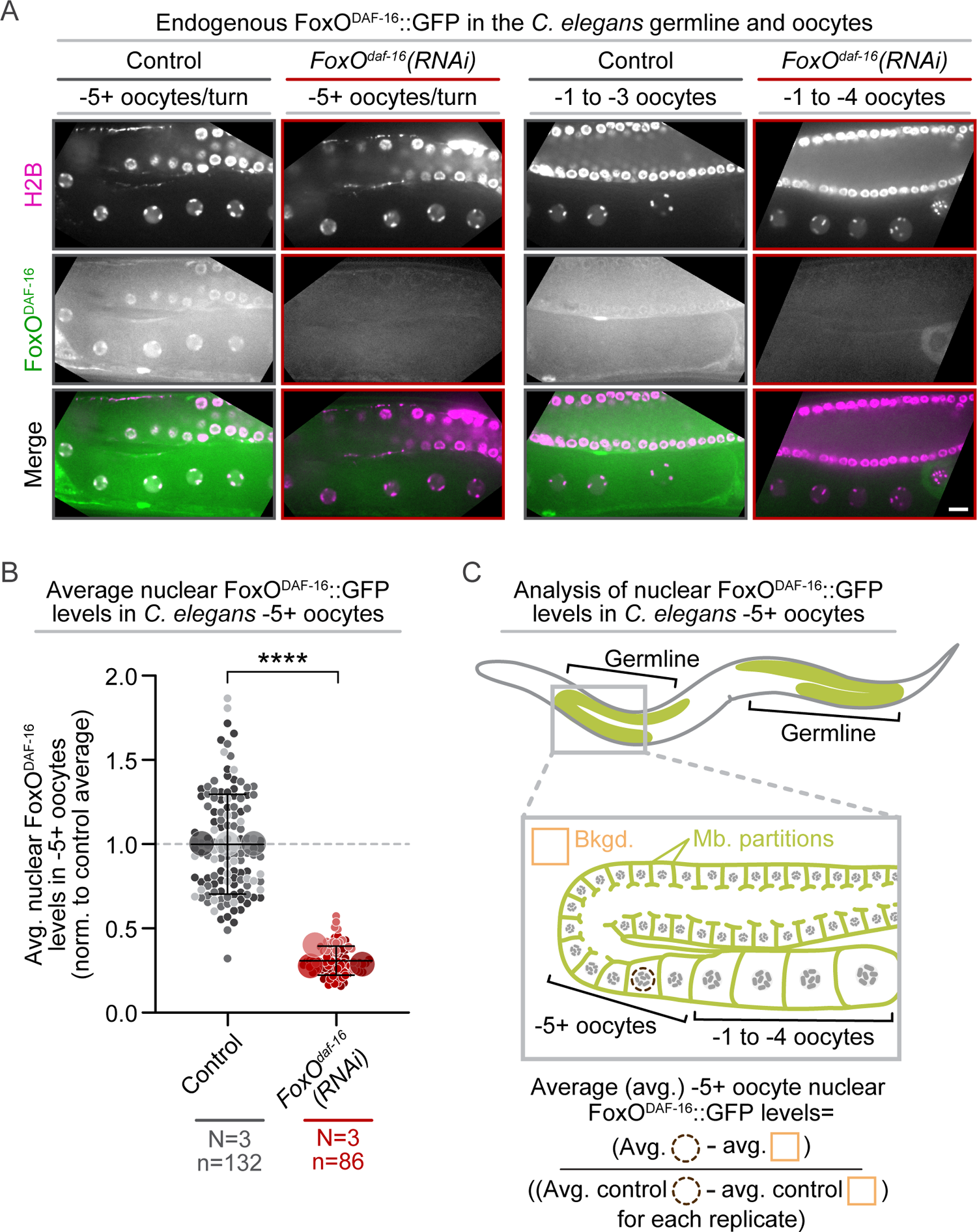
*FoxO^daf-16^(RNAi)* greatly reduces FoxO^DAF-16^ levels in the germline and oocytes. A) Representative single plane images showing FoxO^DAF-16^::GFP (green in merged images) and mCherry::H2B (magenta in merged images) localization in the *C. elegans* germline and oocytes with and without *FoxO^daf-16^(RNAi)*; black box behind some images for display; scale bar=10 μm. **B)** Graph plotting the average (avg.) nuclear levels of endogenously-tagged FoxO^DAF-16^::GFP in control (grays) and *FoxO^daf-16^(RNAi)* (reds) in −5+ oocytes normalized (norm.) to the average nuclear levels in controls. Small circles indicate the average levels in individual −5+ oocyte nuclei and large circles indicate replicate averages; shading indicates results from the same replicate (*e.g.*, lightest shades in both control and RNAi conditions same replicate). Error bars=SD; N=number of experimental replicates; n=number of nuclei scored for genotype by color; ****=p-value ≤0.0001 (Student’s t-test, unpaired; see **Table S1**). **C)** Schematic depicting analysis shown in **(B)** performed on single plane images to measure nuclear FoxO^DAF-16^::GFP levels in −5+ oocytes (dashed brown circle) and average extracellular background signal (orange box); Mb. partitions=plasma membrane partitions (pea green); chromosomes/nuclei (dark/light gray); see also Methods.

**Figure S2:**
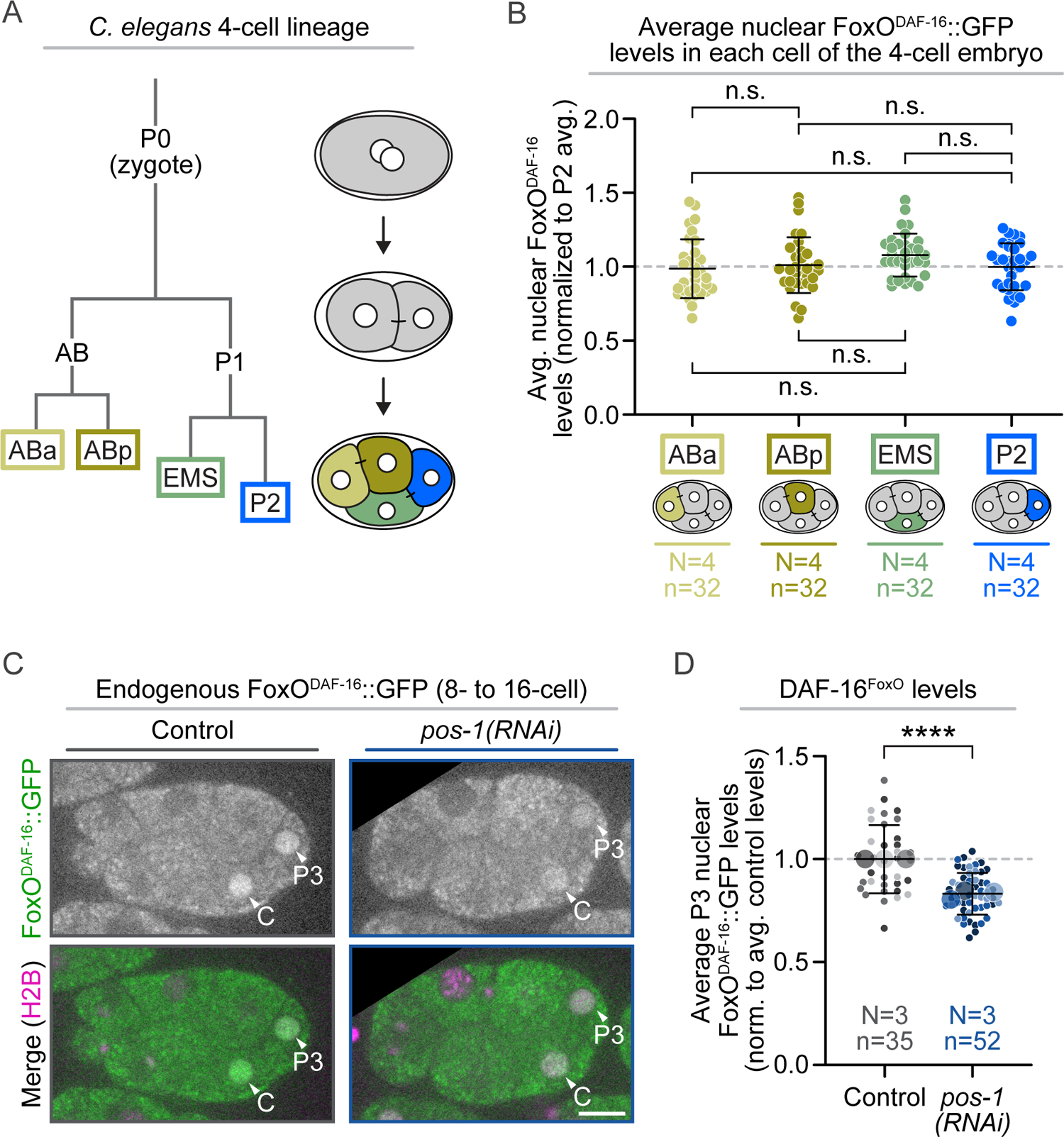
FoxO^DAF-16^ levels are cell type-independent in 4-cell embryos and germ fate dependent in 8-cell embryos. **A)** Schematic depicting the 4-cell *C. elegans* embryo lineage and each cell identity. **B)** Graph plotting the average (avg.) nuclear levels of FoxO^DAF-16^::GFP in the ABa (wheat), ABp (olive green), EMS (sage green), and P2 (blue) cells normalized (norm.) to the average (avg.) nuclear levels in the P2 cell. See **Figure 1B** for representative image. Error bars=SD; N=number of experimental replicates; n=number of nuclei scored for each cell type by color; n.s.=p-value not significant (1-way ANOVA; see **Table S1**). **C)** Representative single plane images showing FoxO^DAF-16^::GFP (green in merged images) and mCherry::H2B (magenta in merged images) localization in 8- to 16-cell embryos with and without *pos-1(RNAi)*; black box behind some images for display; scale bar=10 μm. **D)** Graph plotting the average P3 nuclear levels of FoxO^DAF-16^::GFP in control (grays) and *pos-1(RNAi)* (blues) embryos. Results are normalized to the average nuclear levels in control embryos. Small circles indicate the average levels in individual P3 nuclei and large circles indicate replicate averages; shading indicates results from same replicate (*e.g.*, lightest shades in both control and RNAi conditions from the same replicate). Error bars=SD; N=number of experimental replicates; n=number of P3 nuclei scored for each genotype by color; ****=p-value ≤0.0001 (Student’s t-test, unpaired; see **Table S1**).

**Figure S3:**
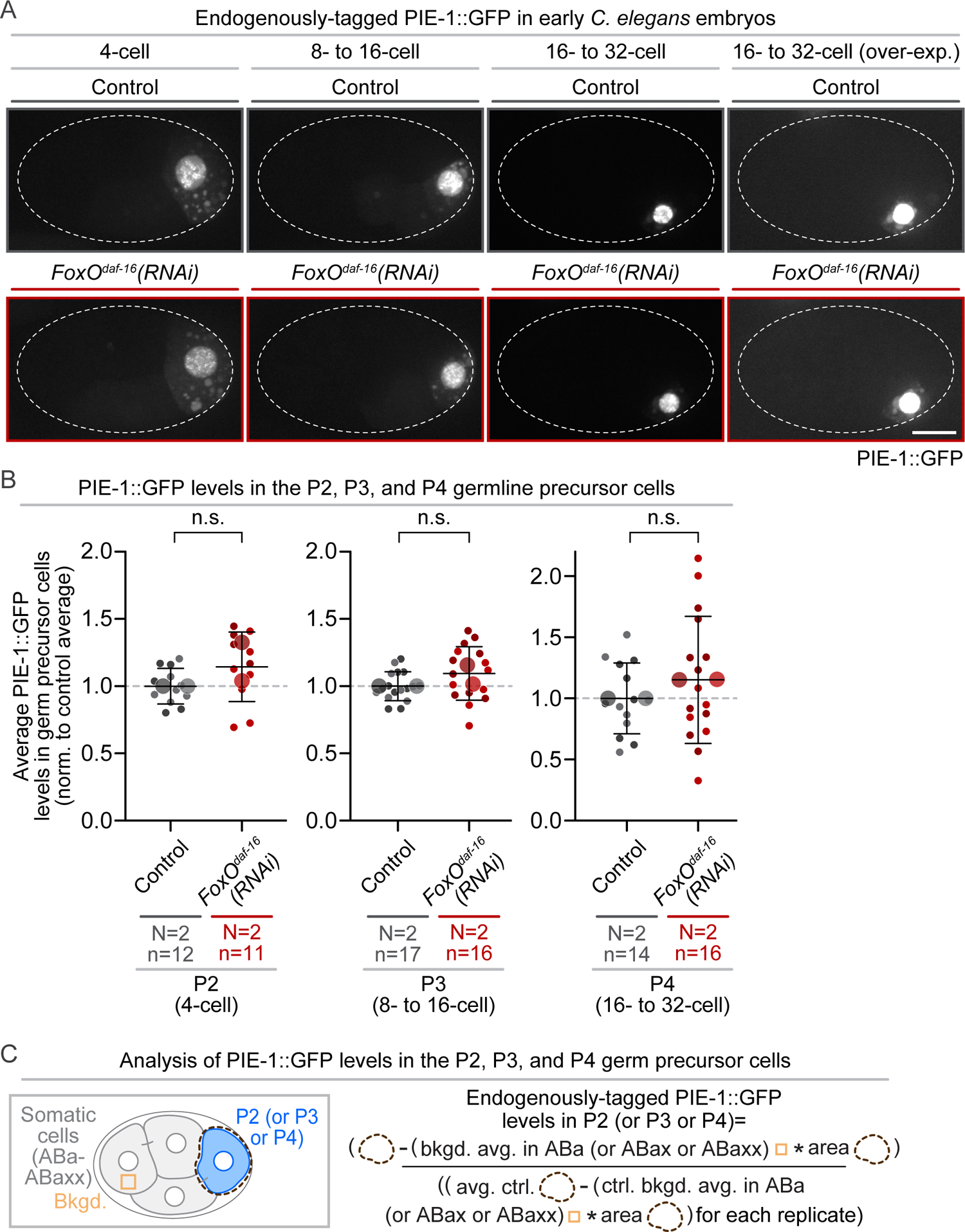
FoxO^DAF-16^ does not regulate the levels of PIE-1 in germ precursor cells. **A)** Representative maximum projection images showing endogenously-tagged PIE::GFP localization in the P2, P3, and P4 germ precursor cells (4-cell, 8- to 16-cell, 16- to 32-cell, and over-exposed (to visualize cytoplasmic p-granules) 16- to 32-cell stage embryos respectively) with and without *FoxO^daf-16^(RNAi)*; scale bar=10 μm. **B)** Graph plotting the average levels of PIE::GFP in control (grays) and *FoxO^daf-16^(RNAi)* (reds) in the P2 (left), P3 (center), and P4 (right) germ precursor cells with and without *FoxO^daf-16^(RNAi)* normalized (norm.) to the average (avg.) levels in control (ctrl.) embryos. Small circles indicate the average levels in individual germ precursor cells and large circles indicate replicate averages; shading indicates results from the same replicate (*e.g.*, lightest shades in both control and RNAi conditions from same replicate). Error bars=SD; N=number of experimental replicates; n=number of germ precursor cells scored for each genotype by color; n.s.=p-value not significant (>0.05) (Student’s t-test, unpaired; see **Table S1**). **C)** Schematic depicting analysis shown in **(B)** performed on sum projected images to measure total PIE-1::GFP levels in individual germ precursor cells and average intracellular background signal in an ABa, ABax, or ABaxx blastomere (orange box); see also Methods.

**Figure S4:**
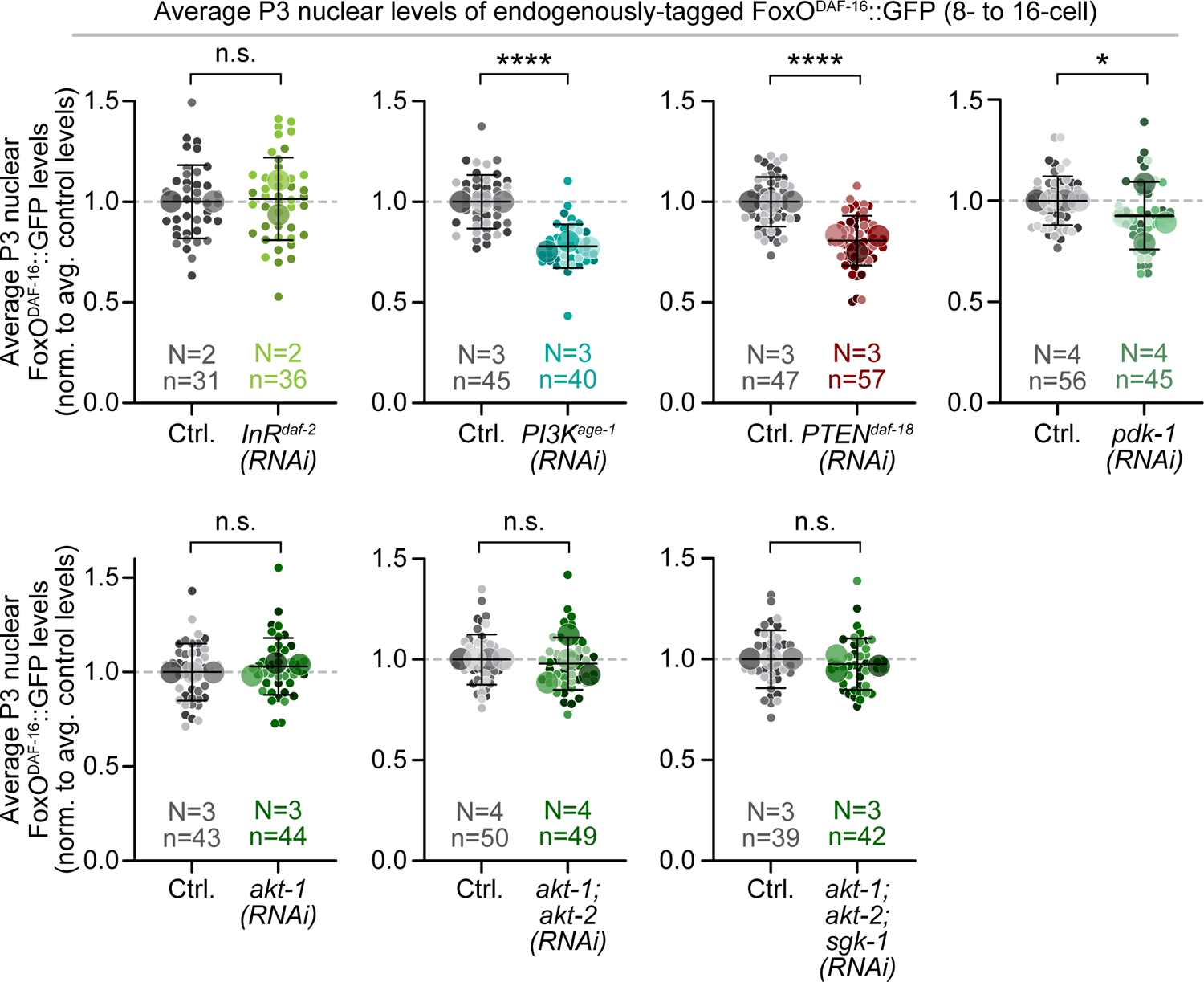
PI3K^AGE-1^ and PTEN^DAF-18^ are required for robust FoxO^DAF-16^ nuclear enrichment in the P3 germ precursor cell. Individual graphs with separated controls for each replicate after RNAi-mediated knockdown of insulin/insulin-like growth factor signaling (IIS) pathway components (same results shown with the controls combined in Figure 3B-C; for schematic of IIS pathway see Figure 1A) normalized (norm.) to the average (avg.) levels in individual replicate control embryos. Graphs plotting P3 nuclear levels of FoxO^DAF-16^::GFP in individual replicate control embryos (left side of all graphs, grays), *InR^daf-2^(RNAi)* embryos (top row, far left; lime greens), *PI3K^age-1^(RNAi)* embryos (top row, center left; teals), *PTEN^daf-18^(RNAi)* embryos (top row, center right; maroons), *pdk-1(RNAi)* embryos (top row, far right; sage greens), *akt-1(RNAi)* embryos (bottom row, left; forest greens), *akt-1(RNAi); akt-2(RNAi)* embryos (bottom row, center; forest greens), *akt-1(RNAi); akt-2(RNAi); and sgk-1(RNAi)* embryos (bottom row, right; forest greens. Small circles indicate the average levels in individual P3 nuclei and large circles indicate replicate averages; shading indicates results from the same replicate (*e.g.*, lightest shades in both control and RNAi conditions from same replicate). Error bars=SD; N=number of experimental replicates; n=number of P3 cells for each genotype by color; n.s.=p-value not significant (>0.05), *=p-value ≤0.05, ****=p-value ≤0.0001 (Student’s t-test, unpaired; see Table S1).

**Figure S5:**
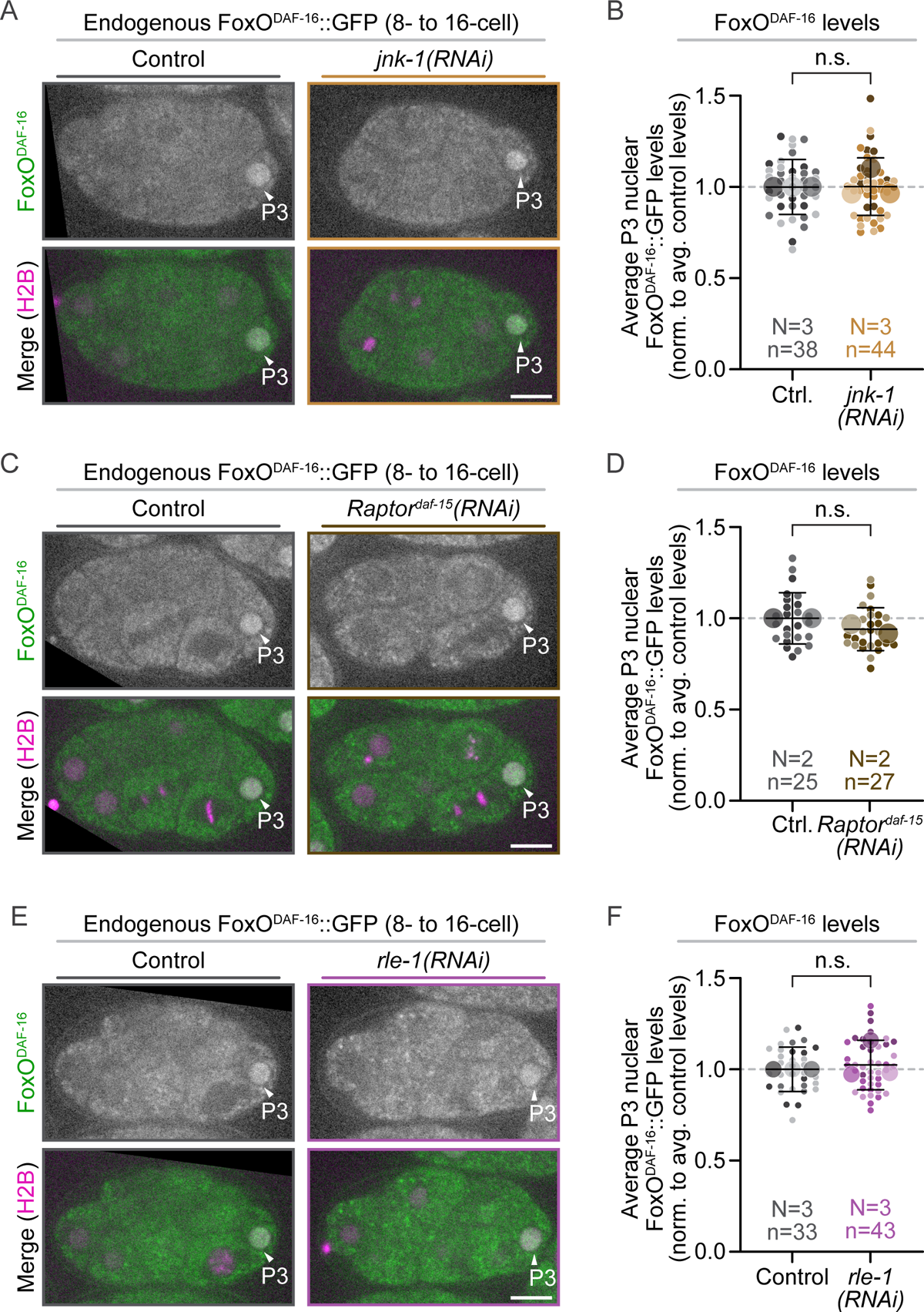
FoxO^DAF-16^ nuclear enrichment in the P3 germ precursor cell is RLE-1, JNK-1, and Raptor^DAF-15^ independent. **A)** Representative single plane images showing DAF-16^FoxO^::GFP (green in merged images) and mCherry::H2B (magenta in merged images) localization in 8- to 16-cell embryos with and without *jnk-1(RNAi)*; scale bar=10 μm. **B)** Graph plotting the average P3 nuclear levels of DAF-16^FoxO^::GFP in control (grays) and *jnk-1(RNAi)* (browns) embryos. **C)** Representative single plane images showing DAF-16^FoxO^::GFP (green in merged images) and mCherry::histoneH2B (magenta in merged images) localization in 8- to 16-cell embryos with and without *Raptor^daf-15^(RNAi)*; scale bar=10 μm. **D)** Graph plotting the average P3 nuclear levels of FoxO^DAF-16^::GFP in control (grays) and *Raptor^daf-15^(RNAi)* (browns) embryos. **E)** Representative single plane images showing FoxO^DAF-16^::GFP (green in merged images) and mCherry::H2B (magenta in merged images) localization in 8- to 16-cell embryos with and without *rle-1(RNAi)*; scale bar=10 μm. **F)** Graph plotting the average P3 nuclear levels of FoxO^DAF-16^::GFP in control (grays) and *rle-1(RNAi)* (purples) embryos. **A, C, E)** Black box behind some images for display. **B, D, F**) Results are normalized (norm.) to the average (avg.) nuclear levels in control embryos for each replicate. Small circles indicate the average levels in individual P3 nuclei and large circles indicate replicate averages; shading indicates results from the same replicate (*e.g.*, lightest shades in both control and RNAi conditions from same replicate). Error bars=SD; N=number of experimental replicates; n=number of cells scored for each genotype by color; n.s.=p-value not significant (>0.05) (Student’s t-test, unpaired; see **Table S1**).

**Figure S6:**
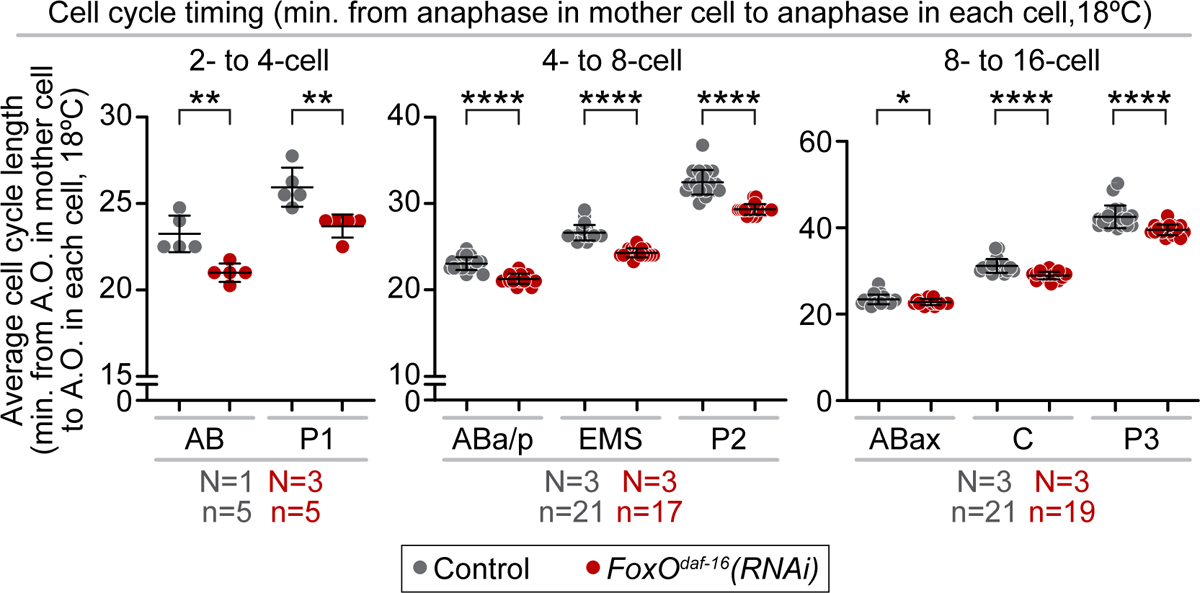
FoxO^DAF-16^ regulates cell cycle timing of early cleavage divisions. Graph plotting the average cell cycle timing in minutes at 18°C (anaphase onset (A.O.) in mother cell to anaphase onset in the indicated cell, for individual cells in control (gray) and *FoxO^daf-16^(RNAi)* (red) embryos. Results are shown for the 2- to 4-cell embryo (AB to P1; far left graph), 4- to 8-cell (ABa/p, EMS, and P2; center graph), 8- to 16-cell embryo (ABax, C, and P3; far right graph). Error bars=SD; N=number of experimental replicates; n=number of cells scored for each genotype by color; *=p-value ≤0.05, **=p-value ≤0.01, and ****=p-value ≤0.0001 (Student’s t-test, unpaired; see **Table S1**).

## Notes

### Competing Interest Statement

The authors have declared no competing interest.

